# Machine Learning of *Pseudomonas aeruginosa* transcriptomes identifies independently modulated sets of genes associated with known transcriptional regulators

**DOI:** 10.1101/2021.07.28.454220

**Authors:** Akanksha Rajput, Hannah Tsunemoto, Anand V. Sastry, Richard Szubin, Kevin Rychel, Joseph Sugie, Joe Pogliano, Bernhard O. Palsson

**Affiliations:** Department of Bioengineering, University of California, San Diego, La Jolla, USA; Division of Biological Sciences, University of California San Diego, La Jolla, CA 92093, USA; Department of Pediatrics, University of California, San Diego, La Jolla, CA, USA; Center for Microbiome Innovation, University of California San Diego, La Jolla, CA 92093, USA; Novo Nordisk Foundation Center for Biosustainability, Technical University of Denmark, Kemitorvet, Building 220, 2800 Kongens, Lyngby, Denmark

## Abstract

The transcriptional regulatory network (TRN) of *Pseudomonas aeruginosa* plays a critical role in coordinating numerous cellular processes. We extracted and quality controlled all publicly available RNA-sequencing datasets for *P. aeruginosa* to find 281 high-quality transcriptomes. We produced 83 new RNAseq data sets under critical conditions to generate a comprehensive compendium of 364 transcriptomes. We used this compendium to reconstruct the TRN of *P. aeruginos*a using independent component analysis (ICA). We identified 104 independently modulated sets of genes (called iModulons), among which 81 (78%) reflect the effects of known transcriptional regulators. We show that iModulons: 1) play an important role in defining the genomic boundaries of biosynthetic gene clusters (BGCs); 2) show increased expression of the BGCs and associated secretion systems in conditions that emulate cystic fibrosis (CF); 3) show the presence of a novel BGC named RiPP (bacteriocin producer) which might have a role in worsening CF outcomes; 4) exhibit the interplay of amino acid metabolism regulation and central metabolism across carbon sources, and 5) clustered according to their activity changes to define iron and sulfur stimulons. Finally, we compare the iModulons of *P. aeruginosa* with those of *E. coli* to observe conserved regulons across two gram negative species. This comprehensive TRN framework covers almost every aspect of the transcriptional regulatory machinery in *P. aeruginosa*, and thus could prove foundational for future research of its physiological functions.

## Introduction

*Pseudomonas aeruginosa* is an opportunistic pathogen that is found in such diverse environments as hospitals, soil, water, and plants. It is one of the major causative agents of hospital-acquired nosocomial infections and a major cause of lung infection in people with cystic fibrosis (CF) ^1^. Indeed, the mortality rate of *P. aeruginosa* infections can reach up to 50% among patients hospitalized with CF and cancer.

All major biological processes in *P. aeruginosa* are controlled by a complex transcriptional regulatory network (TRN) that is yet to be fully elucidated. TRNs constitute the underlying framework for understanding the developmental and physiological responses of organisms ^2^. TRNs, which generally contain the relationships between transcription factors (TFs) and their target genes, play a vital role in programming the bacterial response to diverse stimuli ^3^. Knowledge of the TRN of *P. aeruginosa* and other pathogenic bacteria would be beneficial in elucidating novel drug targets and understanding the functions of their various virulence factors ^2^. Thus, elucidation of these regulatory mechanisms would be beneficial to designing new or combinatorial therapies against *P. aeruginosa* infections. Today, machine learning approaches can be used to establish TRN structure in bacteria if there is sufficient transcriptomic data available for analysis ^4^.

Independent component analysis (ICA), a method to identify independent signals in complex data sets ^5^, has been applied to data sets of bacterial transcriptomes to identify independently modulated sets of genes, called iModulons, and the transcriptional regulators that control them ^6–8^. iModulons have been used to study the adaptive evolution trade-off during oxidative stress under naphthoquinone-based aerobic respiration ^9^, mutations in the OxyR transcription factor and regulation of the ROS response ^10^, and the host response to expression of heterologous proteins ^11^. We have also used ICA to elucidate the TRN structures of *Escherichia coli ^7^*, *Staphylococcus aureus ^6^*, and *Bacillus subtilis ^8^*, which are presented in interactive dashboards on the iModulonDB.org website^12^.

In this study, we apply ICA to high-quality RNA-seq expression profiles of *P. aeruginosa* to decipher the overall structure of its TRN. A previous study used the ADAGE method ^13^ to extract the microbe-host interaction in *P. aeruginosa*, but there is no study available that reveals the structure of its TRN. We incorporated RNA-seq data from diverse conditions such as osmotic stress, low pH, oxidative stress, and micronutrients, and integrated all publicly available data of sufficient quality from the NCBI Sequence Read Archive as of October 20, 2020 ^14^. We assembled the largest possible RNA-seq compendium for *P. aeruginosa*, composed of 364 transcriptomes, and use ICA to reveal the relationship between iModulon activities and specific stimuli. Further, our study identifies several hypotheses from the transcriptomic data that are relevant to *Pseudomonas* infections. Specifically, we characterize the role of many biosynthetic gene clusters (BGCs) which might play a significant role in CF virulence. The TRN structure established here represents a significant advance toward understanding the complex transcriptional regulation of *P. aeruginosa* under different growth conditions.

## Results

### The iModulon structure of *Pseudomonas aeruginosa’s* transcriptome

We assembled the largest possible set of high quality RNAseq profiles for *P. aeruginosa* from the literature and databases and augmented it with new RNAseq profiles for specific conditions of interest. We included RNA-seq expression profiles from two strains of *P. aeruginosa* in this study: PAO1 and K2733 (PAO1*ΔmexB*). The dataset included a range of growth conditions, including micronutrient supplementation, nutrition source variation, osmotic stress, iron starvation, and gene knock-outs (**Supplementary Figure S1a and S1c**). Expression profiles were generated as a part of this study and downloaded from the NCBI SRA database.

After filtering the profiles based on quality criteria (see Methods), we compiled a transcriptomic compendium containing 364 samples (83 new + 281 public expression profiles) (**Figure 1d** and **Supplementary Figure S1c**). All the samples were shown to have Pearson’s correlation coefficient (PCC) of 0.97 between replicates ^4^. To eliminate batch effects, each individual experiment was normalized to a reference condition prior to calculating the iModulons ^4^.

**Figure 1.**
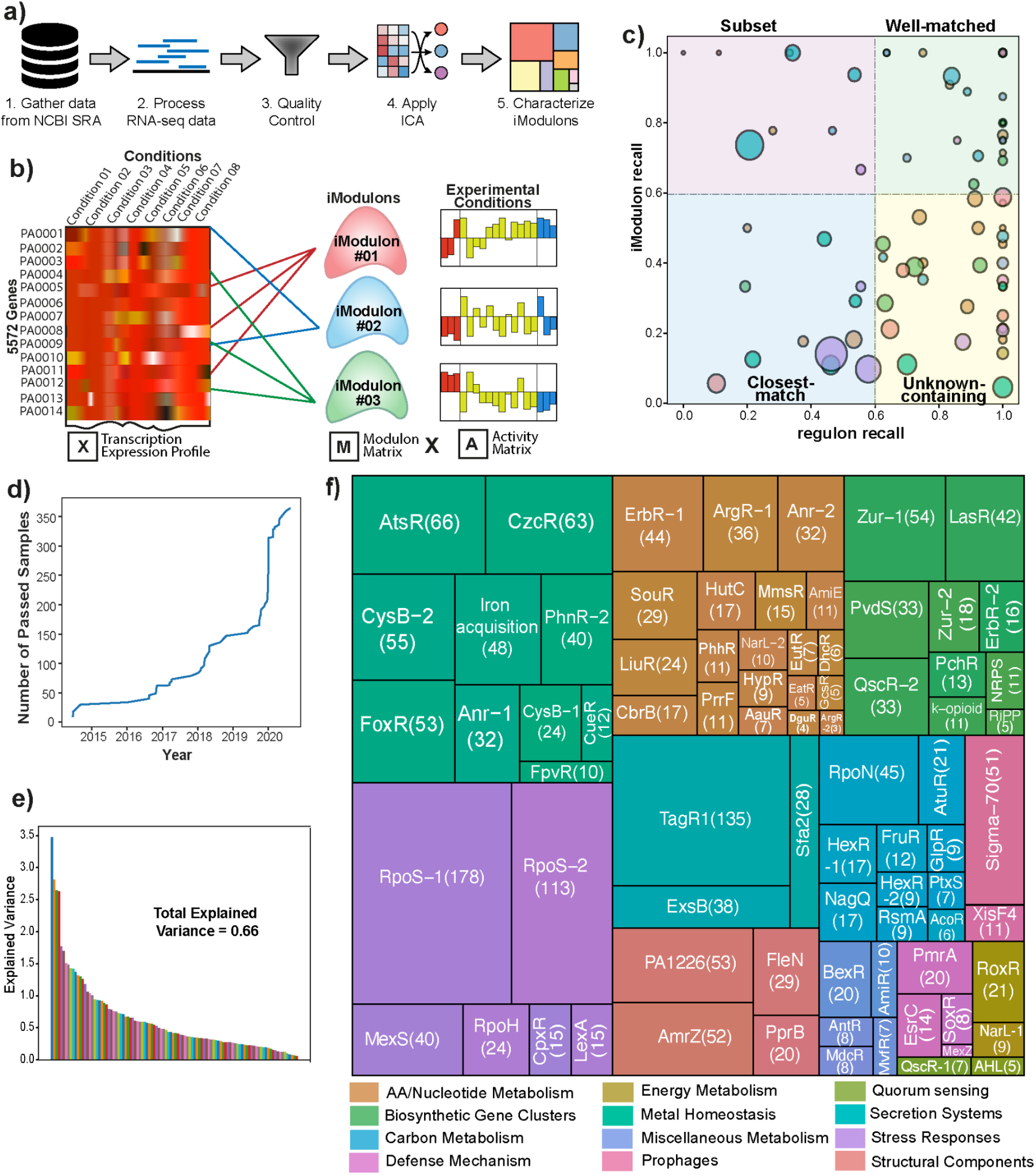
iModulons computed from the Pseudomonas aeruginosa transcriptomic data compendium. a) Overview of the methodology used in the study. It includes gathering high-quality data from the NCBI-SRA as well as generated in the lab. The RNAseq reads were processed and quality control was done. Further, the independent component analysis (ICA) was applied to generate the iModulons that were characterized to get the regulatory networks of P. aeruginosa (Adapted from Sastry et al. ^4^). b) ICA calculates the independently modulated sets of genes (iModulons). A compendium of expression profiles (X) is decomposed into two matrices: the independent components composed of a set of genes, represented as columns in the matrix **M**, and their condition-specific activities (**A**). c) Scatter plot showing the regulon recall versus iModulon recall for all 104 iModulons found in the P. aeruginosa dataset. The scatter plot is divided into four quadrants: Upper right represents the well-matched iModulons; upper left shows iModulons representing a subset of regulon genes; lower right depicts the iModulons containing uncharacterized genes; lower left contains the iModulons with the closest match. d) Plot showing the amount of passed samples per year which is used in the study. e) Bar plot showing the explained variance in all the iModulons with overall explained variance of 0.66. f) Treemap of the 104 P. aeruginosa iModulons. The size of each box represents the fraction of global expression variance that is explained by the iModulon. iModulons are grouped into 12 different categories: AA/Nucleotide Metabolism, Biosynthetic Gene Clusters, Carbon Metabolism, Defense Mechanism, Energy Metabolism, Metal Homeostasis, Miscellaneous Metabolism, Prophages, Quorum sensing, Secretion systems, Stress Responses, and Structural Components. Abbreviations: AA - amino acids

iModulons represent a data-driven, top-down reconstruction of transcriptional regulatory networks and can be characterized using transcriptional regulator binding^7^. To assign transcriptional regulators to iModulons, we compared each iModulon against regulons published in the literature. We compiled a TRN scaffold, using RegPrecise^15^, a manually curated database containing 58 regulons. We then manually searched the literature for additional high-quality transcription factor binding sites. In total, the resulting TRN scaffold contained binding information for 134 TFs and the corresponding regulons. This data is available in **Supplementary Table S1**.

We applied ICA to the transcriptomic compendium to identify independent signals in the data set that represent the effects of transcriptional regulators (**Figure 1a** and **1b**) ^7^. ICA resulted in the identification of 104 iModulons. To annotate each iModulon, their genes were compared with the those in the 134 regulons (**Table S1**) to find statistically significant enrichments (See Methods). For iModulons with strong associations to known regulons, we used ‘iModulon recall’ and ‘regulon recall’ to evaluate our confidence in the associations (**Supplementary Figure S1b**). The iModulon recall represents the fraction of shared genes and the genes in an iModulon while regulon recall is the fraction of shared genes and the genes in a regulon (**Supplementary Figure S1b**).

The relationship between the 134 regulons and the 104 iModulons are grouped into four categories (**Figure 1c)**: 1) The well-matched group (upper right quadrant) includes iModulons with regulon recall and iModulon recall greater than 0.60. The genes in these iModulons correspond to the existing regulons. 2) The subset of iModulons in the upper left quadrant has high iModulon recall and low regulon recall, thus these iModulons represent only a part of a defined regulon. 3) The unknown-containing iModulons (lower right quadrant) have high regulon recall and low iModulon recall. The iModulons in this category mostly include uncharacterized genes. 4) The iModulons with closest match category (lower left quadrant) have low regulon and iModulon recall. These iModulons contain co-expressed genes that showed statistically significant enrichment levels and appropriate activity profiles ^8^. Thus, we identified 104 iModulons that are the regulated gene sets from complementary bottom-up and top-down methods

### Functional classification of the iModulons, their coverage of genes, and how they form the variation in the RNAseq compendium

The 104 iModulons identified were annotated with different functions such as BGCs, secretion systems, stress responses, prophages, metal homeostasis, structural components, amino acid metabolism, and carbon metabolism. We identified 11 iModulons related to BGCs and 14 iModulons related to metal homeostasis. Pathogenicity could be related to three iModulons representing type VI and type III secretion systems ^16^. We also functionally annotated iModulons associated with carbon, amino acids, sulfur, iron, secondary, lipid, and nitrogen metabolism (**Figure 1f**). iModulons associated with transcriptional regulators are functionally annotated using the regulators’ name. As stated above, the regulator’s binding site information was extracted from the RegPrecise database and from various publications (**Supplementary Table S2**).

ICA identified 104 iModulons that represent 65.63% of the variance in the gene expression (**Figure 1e**). Note that the explained variance in the data compendium using iModulons is fundamentally different from the explained variance of other statistical decomposition methods. The iModulons are biologically meaningful signals in the data that are directly connected to knowledge about regulator function. The explained variation in the data set that the iModulons represents is based on molecular mechanistic knowledge about the transcriptional regulators.

Among the 104 iModulons, four contain single genes. The remaining 100 iModulons contain 1835 unique genes. 561 genes were found in more than one iModulon. We have provided the information for each iModulon in the form of an interactive dashboard on iModulonDB.org ^12^. The dashboard is user-friendly and researchers can search or browse the details of iModulons, TRN, genes, or regulators of interest. Such an examination gives 1) a guide to the study of molecular level mechanisms ^17,18^ or 2) systems level mechanisms, such as those of resource allocation through changes in the transcriptome composition between conditions ^7,11^.

### iModulons provide a definition of genomic boundaries of biosynthetic gene clusters

BGCs are clusters of genes that synthesize sercreted secondary metabolites such as pyochelin, pyoverdine, pyocyanin, and bacteriocins^19^. These metabolites are of particular interest because of their range of functions, including antibiotics, anticancer molecules, polymers, nutraceuticals, and many more. The comprehensive antiSMASH software uses sequence comparison to detect BGCs, but assigns BGC borders arbitrarily at 20kb from both ends of the core genes ^20^. BGCs normally form contiguous segments of DNA on the genome. However, iModulons can use co-expression patterns to define the functional gene composition of a BGC, since genes in biosynthetic pathways are usually co-expressed. Thus, iModulons can assist in annotating the BGCs and their accessory functions.

The 104 iModulons contain 11 BGCs out of the 14 predicted BGCs in *P. aeruginosa* using anti-SMASH (**Figure 2a and Supplementary Figure S2a**). The remaining 3 BGCs may not have been transcriptionally activated by the conditions represented in the transcriptomic dataset analyzed and thus ICA cannot detect them. The ErbR-2 iModulon contains coregulated genes which are predicted to be redox-cofactors like Pyrroloquinoline quinone (PQQ) (**Figure 2b and c**). The BGC’s boundaries defined by antiSMASH are arbitrarily marked from PA1977-PA1997. However, the iModulon captured by ICA identifies an independent transcriptional signal from PA1975 to PA1990. This BGC also includes the PA0565 gene which has a potential role in post translational modification (PTM), protein turnover, and chaperones (**Figure 2d**). PTM is an important process during the synthesis of secondary metabolites ^21^. All 11 iModulons related to BGCs can be used to annotate their boundaries (**Supplementary Figure S2b**).

**Figure 2.**
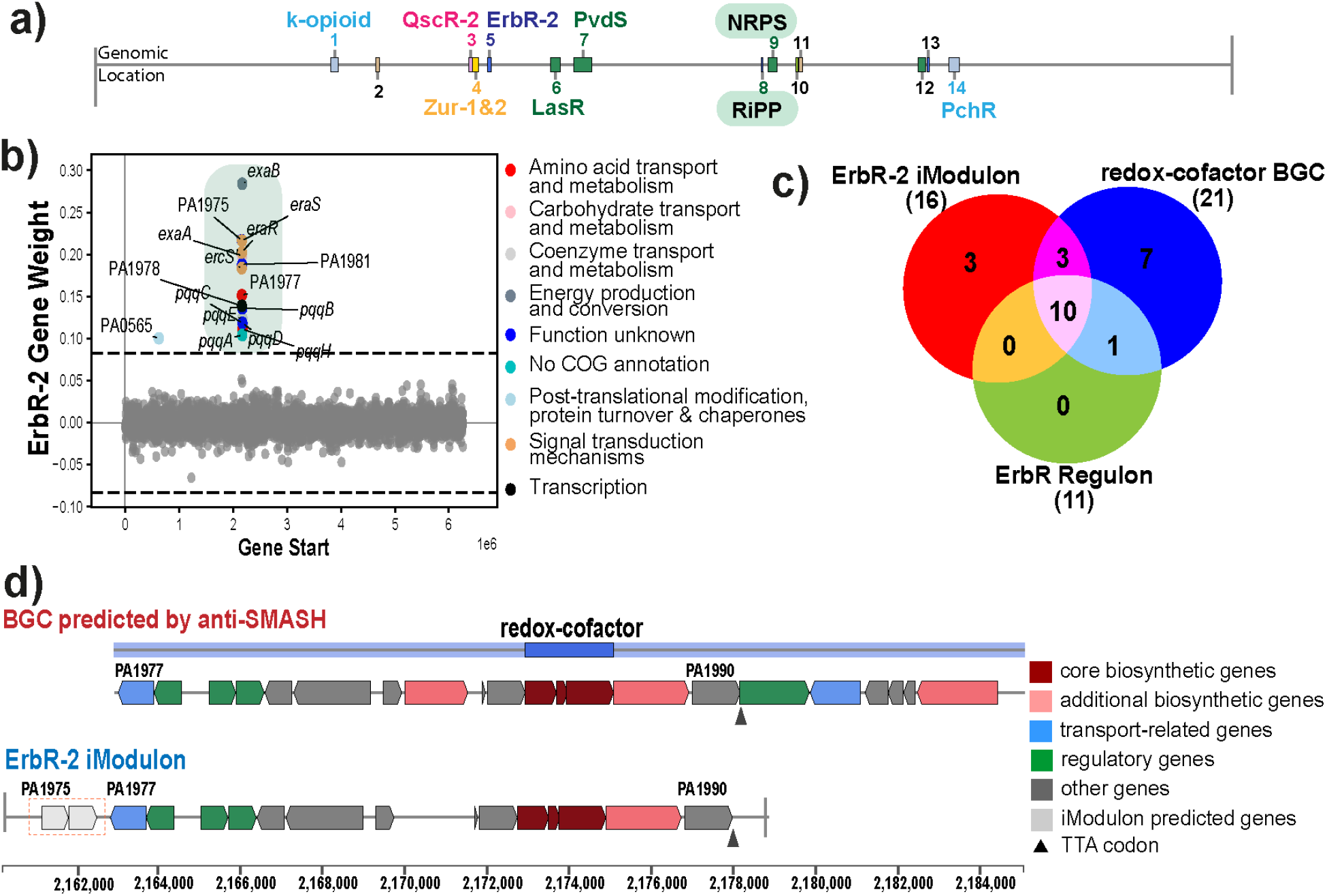
iModulons can aid in the definition of genomic boundaries of biosynthetic gene clusters (BGCs). a) Genomic locations of the 14 predicted BGCs in the P. aeruginosa PAO1 by using the anti-SMASH software. b) Scatter Plot showing the gene weights of the ErbR-2 iModulon with the color depicting the COG categories of the genes that it contains. c) Venn diagram depicting the status of the genes in the ErbR-2 iModulons, ErbR regulon, and the predicted redox-cofactor BGCs by using the anti-SMASH software. d) Genomic overview of the redox-cofactor BGCs predicted by the anti-SMASH software, alongside the iModulons whose boundaries are defined by genes between the PA1975-PA1990.

### iModulons elucidate responses to simulated lung environment in CF patients

*P. aeruginosa* is one of the main bacterial pathogens responsible for causing lung deterioration in CF patients ^22^. The sputum of CF patients shows increased concentrations of copper (Cu), zinc (Zn), iron (Fe), and N-acetyl glucosamine (GlcNAc), compared to the sputum from individuals with healthy lungs. These micronutrients are correlated with increased severity of the disease^23^. iModulons can be used to analyze transcriptomic changes during growth in conditions with altered micronutrients or GlcNAc concentrations.

The NagQ operon is responsible for catabolism of GlcNAc ^24^. *P. aeruginosa* gets GlcNAc from various sources in the host. For example, in the lungs, *P. aeruginosa* obtains GlcNAc by degrading the eubacteria. The structural components (chitin and peptidoglycan) of the degrading eubacteria serve as a major source of GlcNAc ^24^. Thus, the role of GlcNAc in driving *P. aeruginosa* virulence and persistence in human lungs is important and provided an impetus for additional data generation.

We grew *P. aeruginosa* PAO1 in M9 minimal media supplemented with different concentrations of GlcNAc (1, 2, 4, and 8 g/l) as the bacteria’s sole carbon source and examined its impact on iModulons related to BGCs, secretion systems, carbon metabolism, and amino acid metabolism (**Figure 3a-c**). Among the BGCs, we found an increased expression of PvdS, which regulates synthesis of pyoverdine, and a novel cluster named RiPP. The PvdS regulon is responsible for the synthesis of pyoverdine, a siderophore, and led to increased pathogenesis during the lung infections ^22^. The increased expression of pyoverdine in the presence of GlcNAc has been previously reported in the *Streptomyces* species ^25,26^ but not in *P. aeruginosa*. Contrary to PvdS, we found decreased expression of the pyocyanin secreting BGCs (QscR-2 iModulon).

**Figure 3.**
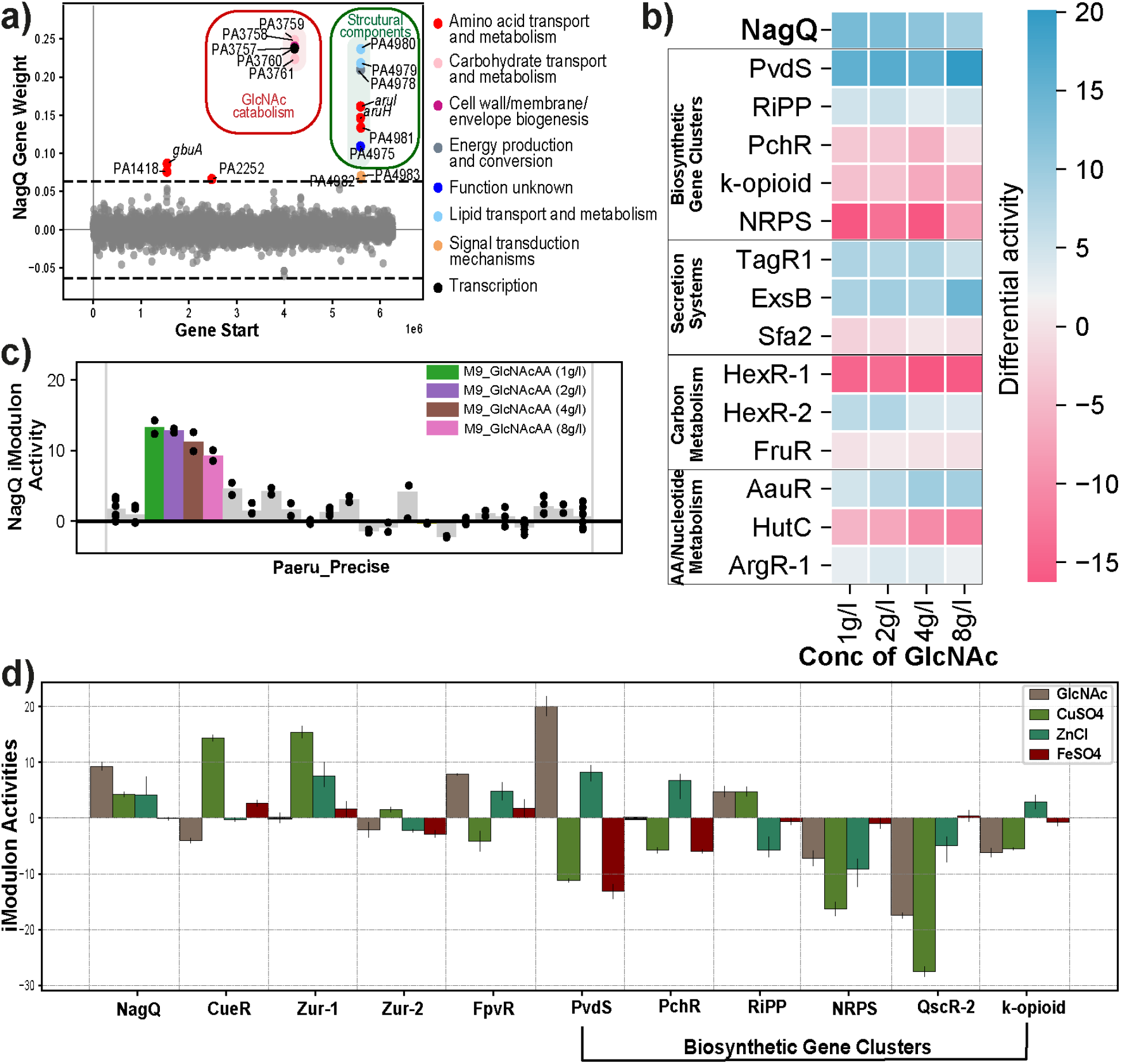
iModulon responses to GlcNAc culture. a) Scatter Plot showing the gene weights of the NagQ iModulon; the color depicts the COG categories. The NagQ iModulons have two regulons; one is GlcNAc catabolism and other is related to structural components. b) Heat map depicting the activity of selected iModulons in different concentrations of GlcNAc (1g/l, 2g/l, 4g/l, and 8g/l). It describes the change in differential activities in NagQ, biosynthetic gene clusters, secretion systems, carbon metabolism, amino acid metabolism, and nucleotide metabolism. c) Activity plot of the conditions expressed in NagQ iModulon in the Paeru_Precise. d) Plot showing iModulon activities in the presence of N-acetyl glucosamine (GlcNAc), ZnCl, CuSO_4_, and FeSO_4_ micronutrients. The iModulons include the micronutrient metabolism (NagQ, CueR, Zur-1, Zur-2, FpvR) and the biosynthetic gene clusters (PvdS, PchR, RiPP, NRPS, QscR-2, and k-opioid).

We also found a previously unannotated BGC, named RiPP (ribosomally synthesized and post-translationally modified peptide), which showed increased expression in the presence of GlcNAc. The novel RiPP contains a DUF692-associated bacteriocin producing domain, as predicted by anti-SMASH (**Supplementary Figure S3a**). This DUF692-associated bacteriocin has not been reported in *P. aeruginosa*; however the evidence of its presence is reported in *Streptomyces* and *Methanobacteria* sps ^27^. Because of its expression in the presence of GlcNAc, this novel BGC of *P. aeruginosa* has a probable role in CF. It was also found to be expressed in the presence of sodium hypochlorite (NaOCl) (**Supplementary Figure S3b**). The role of hypochlorite is also established in the pathogenicity of the CF patients ^28^ and is one of the major oxidants known to be generated by activated phagocytes ^29^.

### iModulons reveal coordinated expression of secretion systems

The role of the secretion systems is well known to increase the pathogenicity of *Pseudomonas* by the secretion of virulence factors ^30^. We found an increased expression of the H1-Type VI secretion system (H1-T6SS) and Type III secretion system (T3SS) in the presence of GlcNAc (**Figure 3b**). The H1-T6SS is known to target the prokaryotic cell and contributes to the survival advantage of *P. aeruginosa*. In the H1-T6SS, the Tse1-Tse3 provides a fitness advantage to *P. aeruginosa* during interspecies communication ^31^. In comparison, the T3SS in *P. aeruginosa* is one of the major virulence factors that contributes to the cytotoxicity and acute infections. T3SS is used to inject the effector proteins into the host cells ^32^. The activation of the secretion systems might be helpful to export the products of the BGCs, such as pyoverdine and RiPP, outside *P. aeruginosa ^32^*. Thus, we found that specific BGCs for pyoverdine and the uncategorized RiPP as well as secretion systems are activated in the presence of GlcNAc, and are potentially linked to the deterioration of health in CF patients during infection with *P. aeruginosa.*

In addition to GlcNAc, the CF lung microenvironment contains elevated levels of Cu, Zn, and Fe ^23^. We have generated the transcriptomic profiles for Cu, Zn, GlcNAc, Fe; and examined the iModulon activities in the presence of these metals (**Figure 3d**). We found that the CueR iModulon is upregulated in the presence of Cu. The expression of the Zur-2 and FpvR iModulons are repressed in the presence of Zn and Fe, respectively. Both Zur-2 and FpvR function in concert with other proteins to bring in Zn and Fe, respectively, into the cells during growth in conditions with low Zn or Fe ^33,34^. However, the Zur-1 iModulon shows activation in the presence of Zn because the genes responsible for binding the Zn show downregulation in this iModulon. Interestingly, we also found that iModulons related to the secretion of pyochelin and pyoverdine, as well as a novel bacteriocin producing (RiPP), are upregulated in presence of these micronutrients. The PchR and PvdS iModulons are responsible for the expression and secretion of the siderophores pyochelin and pyoverdine, respectively. The PchR iModulon showed increased activity in the presence of Zn, while the PvdS is activated during growth with both GlcNAc and Zn.

### iModulons describe central metabolic pathways

We found multiple iModulons related to central carbon metabolism, such as those that are involved in the Entner–Doudoroff, peripheral, and glycolytic pathways (**Supplementary Figure S4**). Among the identified metabolic iModulons, NagQ is related to the catabolism of GlcNAc into glucosamine-6-phosphate (GlcN-6P) and then to Fructose-6-phosphate, a key glycolytic intermediate. As mentioned previously, GlcNAc is well known for its role in the pathogenicity in CF patients by activating various BGCs and secretion systems to target other bacterial species as well as host cells ^24^. During growth in lungs with CF, the catabolism of GlcNAc serves as the main carbon and energy source for *P. aeruginosa ^35^*. Furthermore, iModulons related to the catabolism of the ethanolamine, glycerol, fructose, and 2-ketogluconate describe the state of the metabolic network when these substrates serve as carbon sources (**Figure 4a, Supplementary Figure S4 and S5a)**.

**Figure 4.**
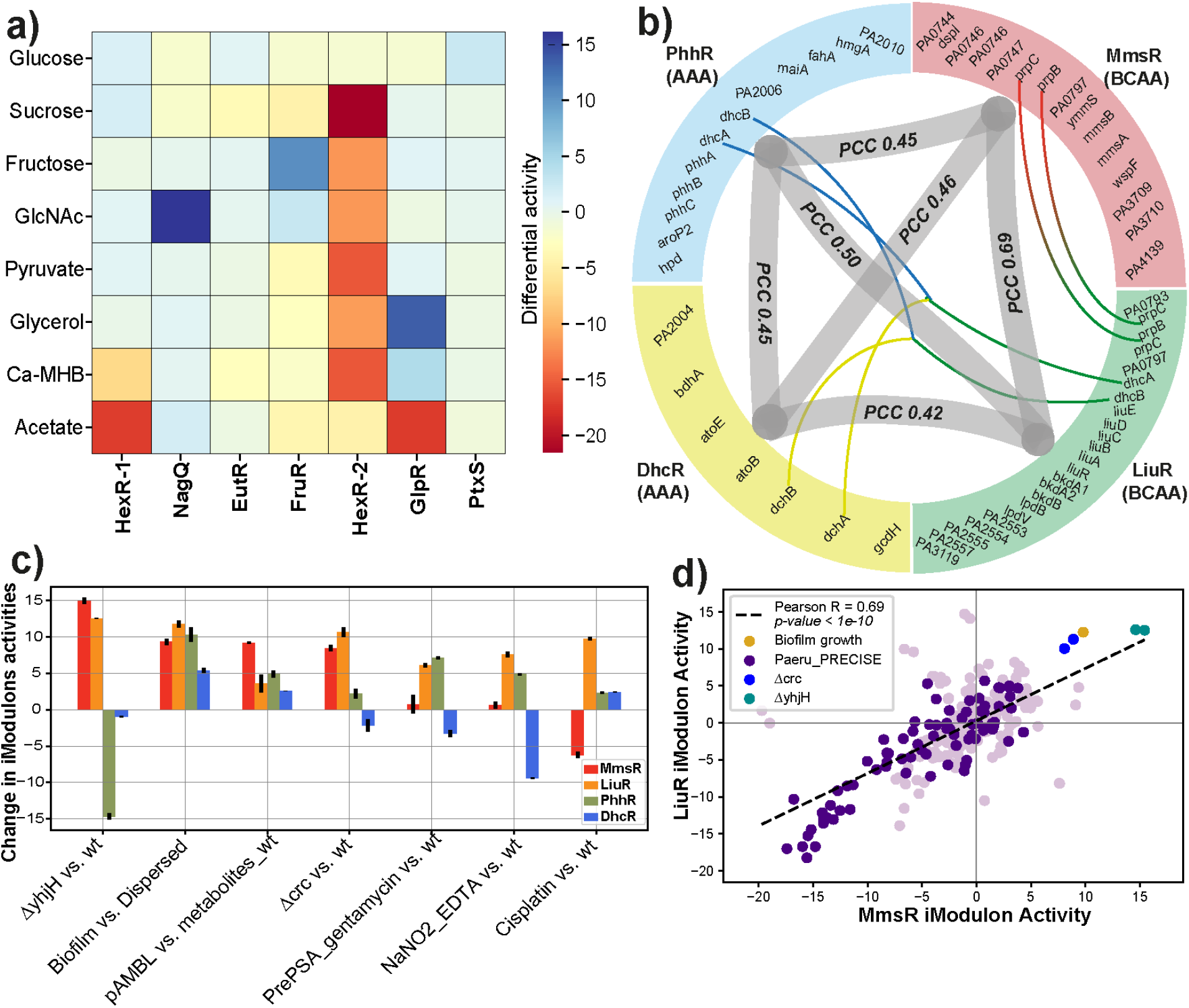
iModulons related to Carbon metabolism and Amino acid/Nucleotide metabolism. a) Heat map depicting the differential activity of glucose, sucrose, fructose, N-acetylglucosamine, pyruvate, glycerol, Ca-MHB, and acetate w.r.t. HexR-1, NagQ, EutR, FruR, HexR-2, GlpR, and PtxR iModulons. b) Correlation plot among the BCAA (LiuR and MmsR) and the AAA (DhcR and PhhR). The outer layer is divided into the four arcs which depict the four different iModulons. Thin lines represent the common genes among the iModulons, and the thick line connecting different iModulons depicts the Pearson correlation coefficients (PCC). c) Bar plot representing the iModulon activities of MmsR, LiuR, and PhhR under different conditions. The x-label shows some conditions used in the study. The ‘ΔyhjH vs. wt’ is the knockout of the yhjH, ‘Biofilm vs. Dispersed’ is the biofilm mode of growth, ‘pAMBL vs. metabolite_wt’ is the pAMBL plasmid showing overexpression of metabolites, ‘Δcrc vs. wt’ is the deletion of the global regulator of crc, ‘PrePSA_gentamycin vs. wt’ is the pre-PatH-Cap library of P. aeruginosa (“PSA” PAO1-GFP) treated with gentamycin,’NaNO2_EDTA vs. wt’ is the presence of sodium nitrite and EDTA in the media, and ‘Cisplatin vs. wt’ is the presence of cisplatin and bile in the media. d) Scatter plot showing the correlation between the BCAA pathways iModulons, i.e., LiuR and MmsR, with the PCC of 0.69.

Several of the identified iModulons mapped onto amino acid metabolic pathways, such as branched chain amino acids (MmsR, AtuR, PrrF, and LiuR), aromatic amino acids (PhhR and DhcR), arginine catabolism (CbrB), histidine utilization (HutC), arginine succinyltransferase (ArgR-1 & 2), arginine deaminase (ArcR), and L-hydroxyproline (HypR) (**Supplementary Figure S4**). We have found significant correlations among the iModulons regulating the branched-chain amino acid (BCAA) and aromatic amino acid (AAA) pathways (**Figure 4b**). Amino acids provide a supplemental nitrogen source and play a vital role in biofilm formation ^36^, and it has been hypothesized that amino acids promote biofilm production in CF patients ^37^. In support of these hypotheses, our data showed that iModulons related to amino acid metabolism pathways had higher activities during growth in biofilm conditions compared to planktonic growth (**Figure 4c and Supplementary Figure S5b**).

### Phosphodiesterases stimulate the expression of branched-chain amino acids

c-di-GMP is a secondary messenger that regulates various important cellular processes like quorum sensing, biofilm formation, and pathogenicity^38^. YhjH is a c-di-GMP phosphodiesterase and, upon induction, it decreases c-di-GMP levels^39^. A decrease in c-di-GMP levels leads to a decrease in the biofilm formation and increased biofilm dispersal. We found the knockouts of YhjH (ΔyhjH, PRJNA381683)^40^ led to increased expression of BCAA metabolism iModulons, subsequently increasing intermediates of the tricarboxylic acid (TCA) cycle (**Figure 4c**). Likewise, the deletion of Crc (Δ*crc*) also led to increased expression of the BCAA iModulons (**Figure 4d**), similar to YhjH. Crc is a global regulator that represses succinate metabolism and BCAA assimilation in *P. aeruginosa* and *P. putida* ^41^.

From our analysis, we observe that YhjH modulates biofilm formation, as previously reported by Zhang *et al ^42^*.Thus, YhjH may be used as an important target to control the biofilm formation and pathogenicity of *P. aeruginosa*. Activities of the identified iModulons were therefore able to untangle complex relationships between metabolites, transcriptional regulators, and lifestyle in *P. aeruginosa*.

### Correlated activity changes of iModulons lead to definition of Stimulons

We have clustered the iModulons based on their correlation as set (**Figure 5a** and **Supplementary Figure 6a**). Though iModulons are independently modulated throughout the transcriptome, clusters of iModulons may be similarly expressed across most conditions in the compendium and only diverge from one another under select conditions. Thus, a cluster of iModulons with coordinated activity changes can be interpreted as a “stimulon”. Such clusters of iModulons are of interest for understanding the broader structure of transcriptional regulation ^43^. For example, we have found that sulfur stimulon {AtsR, CysB-1, and CysB-2} and iron stimulon {FpvR, PvdS, PchR, FoxR, and Iron acquisition} are among the top clustered stimulons in *P. aeruginosa.*

**Figure 5.**
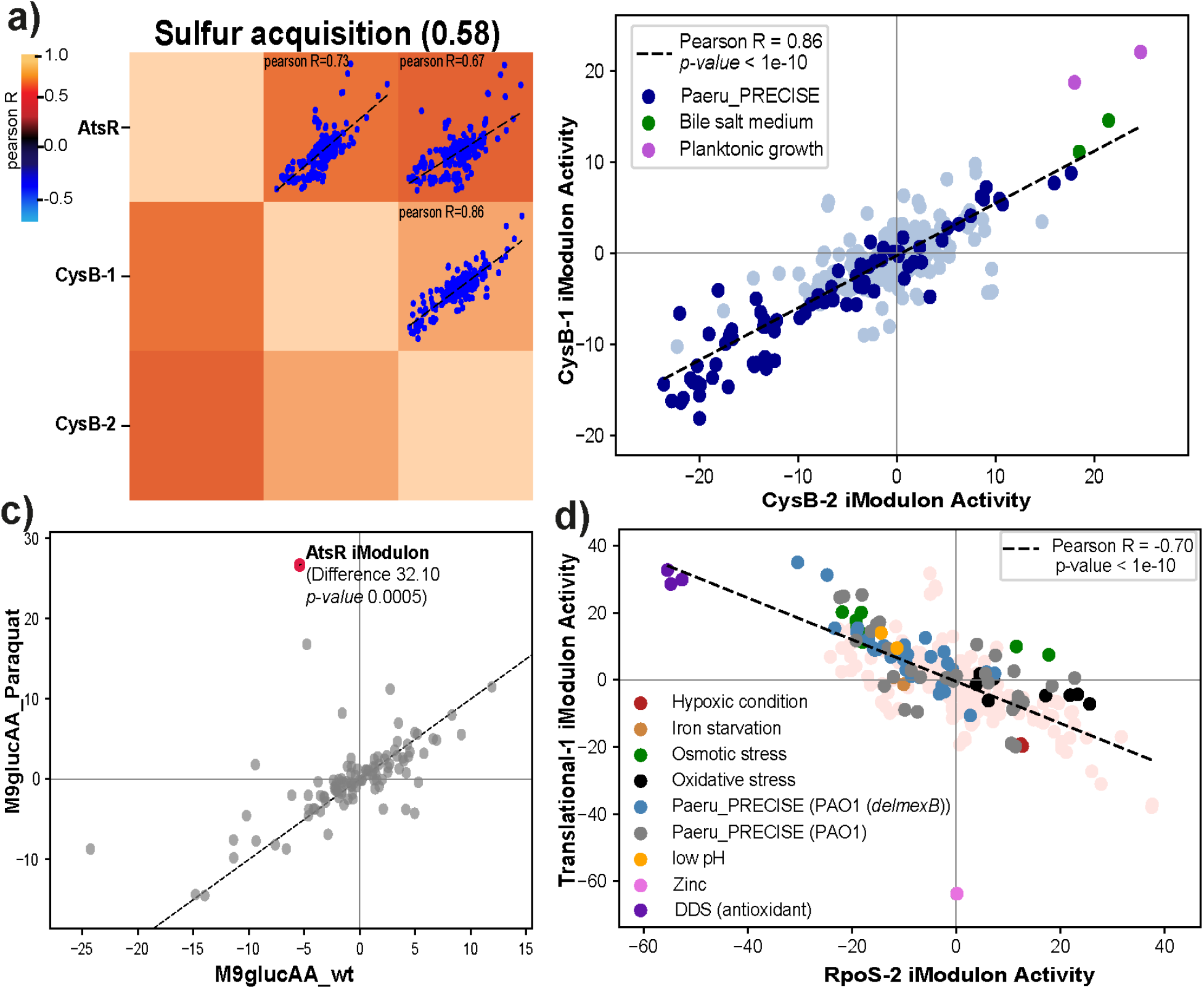
Activity clustering of the iModulons among P. aeruginosa defines stimulons. a) Sulfur acquisition cluster includes the grouping of AtsR, CysB-1, and CysB-2 iModulons with silhouette score of 58. b) The scatter plot showing correlation between the CysB-1 and CysB-2 iModulons with PCC of 0.86. Both the iModulons show high activity in the planktonic condition and bile salt medium of P. aeruginosa. c) Differential iModulon activity (DIMA) plot of the M9glucAA and Paraquat v/s the control condition showing the upregulation of the AtsR iModulon. d) The RpoS-2 iModulon activities were anti-correlated with the Translational-1 iModulon activities. All the stress conditions (hypoxia, iron starvation, osmotic stress, oxidative stress, and low pH) were highlighted with different colors.

#### Sulfur acquisition

The iModulons AtsR, CysB-1, and CysB-2 form a sulfur acquisition stimulon (**Figure 5a**). AtsR is a transcription factor that encodes the ABC transporters of sulfate and other ions. We observed that the AtsR iModulon was activated during oxidative stress (paraquat supplementation) (**Figure 5c**). The relationship between sulfate limitation and the oxidative stress response has been established in *E. coli* ^44^ but not in *P. aeruginosa*. The CysB-1 and CysB-2 regulators modulate sulfur uptake and cysteine biosynthesis, as well as influence the genes involved in host colonization and virulence factor production ^45^. The two CysB iModulons are highly expressed in planktonic growth conditions as well as in the presence of bile (**Figure 5b**). However, CysB’s direct connection with bile has not been previously established in the literature. Taurine, a sulfur-containing amino acid, is one the primary components of bile acids. We hypothesize that *P. aeruginosa* upregulates its sulfur acquisition genes in response to the presence of taurine in the conjugated bile acid. Interestingly, in certain patients with CF, there can be microaspirations of bile into the lungs, and studies have shown that bile affects the transition of *P. aeruginosa* into biofilms ^46^. Therefore, CysB may play an important role in the pathogenicity of *P. aeruginosa* in CF lungs through its role in acquiring sulfur from bile aspirations.

#### Iron acquisition

We identified a cluster of five iron-related iModulons (FpvR, PvdS, PchR, FoxR, and Iron acquisition) (**Supplementary Figure 6a**). The iron stimulon contains a set of five correlated iModulons. The five iModulons involved in this cluster contained genes involved in the uptake of iron through endogenous (pyoverdine, *fpv, pvd*) or exogenous (xenosiderophores, FoxR and heme) carriers ^47^. The activities of both the endogenous PvdS and exogenous FoxR iModulons were upregulated during the presence of the chelator EDTA and during planktonic growth (PCC 0.67) (**Supplementary Figure 6b**). The iron acquisition iModulon was previously uncharacterized and known as Uncharacterized-13, which was further annotated to be involved in iron acquisition by clustering analysis. Additionally, the presence of an uncharacterized iModulon (Uncharacterized-13) in this cluster allowed us to annotate its potential function, which we hypothesize as a role in pyoverdine synthesis. Several genes, such as PA2531, PA4709, *phuR*, *opmQ*, *pvdT*, *pvdR*, *pvdE,* and PA2412 are shared between the PvdS and Uncharacterized-13 iModulons (**Supplementary Figure 6c**). Thus, our analysis provides insight into the interconnectedness of iron acquisition systems in *P. aeruginosa*.

### iModulons show ‘Fear vs. Greed’ Trade-off

In previous studies of *E. coli* and *S. aureus* transcriptional regulation, a trade-off between the expression of translation machinery and stress-hedging genes was observed ^6,7^. This global trade-off was termed the “Fear vs. Greed” trade-off.

The allocation of the resources to the optimal growth (greed) versus its allocation towards the bet-hedging strategies to attenuate its effect of the stressors in the environment (fear) ^7,48^ was demonstrated using two iModulons in *E. coli*. We identified two iModulons in *P. aeruginosa* (Translation-1 and RpoS-2 iModulons) that were orthologous to these *E. coli* iModulons (translation and RpoS iModulons) (**Supplementary Table S3**). The RpoS-2 iModulon includes the sigma factor (RpoS) and a central regulator of the stress response that allows cells to survive environmental challenges. The translational iModulon represents the translational machinery like ribosomal proteins and growth-related function of the transcriptome. We identified an anti-correlation relationship between the RpoS-2 iModulon and the Translational-1 iModulon (**Figure 5d**). Further, the RpoS-2 iModulon also showed correlation (PCC 0.61, *p-value*<10^−10^) with the expression level of the *rpoS* gene, which was also observed in *E. coli* (**Supplementary Figure 6d**).

These results suggest that the ‘Fear vs Greed’ trade-off relationship is conserved among bacterial species.

## Discussion

We have constructed a large compendium of *P. aeruginosa* transcriptomes from all publicly available high-quality data, compiled a TRN of 134 regulons from literature, and computed and characterized a data-driven TRN of 104 iModulons that matches well with the literature. The regulons are based on targeted biomolecular studies, whereas iModulons result from data analysis of a global compendium of transcriptomic data. These complementary approaches to TRN elucidation synergize well. The iModulons were effective for clarifying BGC groups (including identifying a new BGC), characterizing simple and disease-relevant growth conditions from a transcriptomic perspective, clustering functional groups of genes, and comparing regulatory modules across organisms.

From our analysis, we find that iModulons are likely useful in determining the boundary of BGCs. Various initiatives have been undertaken to confirm the boundaries of BGCs, but existing, arbitrary rules do not capture the important feature that they be co-transcribed ^49^. Thus, we present improved annotations for 11 *P. aeruginosa* BGCs. Apart from the other BGCs, we found a novel BGC (the RiPP) that is known to release bacteriocin and to upregulate micronutrients known to be present in high amounts in the lungs of CF patients in the presence of GlcNAc and Cu.

*P. aeruginosa* is an opportunistic pathogen and an important member of the ESKAPEE group of pathogens. It is one of the dominant species responsible for worsening the condition of CF patients ^50^. Interestingly, we have found that BGCs and secretion systems play a significant role in intensifying infections in CF patients. Several BGCs, including the newly discovered RiPP discussed earlier, are upregulated under the GlcNAc and Cu environments representative of CF microenvironments. Apart from the BGCs, we find upregulation of the secretion systems (H1-T6SS and T3SS) in the presence of the GlcNAc. These secretion systems are well known to increase the pathogenicity in the host ^32,51^. Thus, we hypothesize that the BGCs and the secretion system might play an important role in pathogenesis of *P. aeruginosa* during growth in the CF lung. Bernier *et al*, demonstrated that several amino acids promoted biofilm formation of *P. aeruginosa ^37^*; we find that the AAA and the BCAA related iModulons show upregulation in the biofilm mode of growth.

From the functional clustering of iModulons can annotate an uncharacterized iModulon (Uncharacterized-13) which might be responsible for additional iron acquisition. Furthermore, we found a potential correlation of the bile and sulfur acquisition, which might be an important factor for *P. aeruginosa* infection in CF patients. We performed interspecies iModulon comparison. We found 20 iModulons from *P. aeruginosa* showing high correlation with the *E. coli* iModulons, with the translational iModulon being the most important among them. Additionally, we find that the stress related iModulon (RpoS-2) shows anti-correlation with the translational (Translational-1) iModulon, which demonstrates the survival strategy of *P. aeruginosa* under stress conditions.

All the activity and expression profiles as well as the details of iModulons would be very useful for microbiologists to tackle the nosocomial infection causing *P. aeruginosa*. The code for the pipeline is available on github (https://github.com/akanksha-r/modulome_paeru1.0). The framework of the Pseudo Precise iModulons would be helpful to elucidate the regulatory metabolic networks, transcription factors, and various cross-talk among mechanisms. To browse or search dashboards for each iModulon and gene analyzed in this study, visit iModulonDB.org (https://imodulondb.org/dataset.html?organism=p_aeruginosa&dataset=modulome).

In this study, we implemented machine learning to identify the TRN in *P. aeruginosa*. We incorporated high quality transcriptomics data, both from in-house generated data as well as all publicly available, high quality data from the SRA database, to get the independently co-regulated sets of genes (iModulons) which provide a genome-wide, top-down perspective of the TRN of *P. aeruginosa*. We have demonstrated its usefulness for characterizing BGCs, metabolism, and virulence, and its wide scope could enable additional insights into many other processes in *P. aeruginosa*. It may also serve as the basis for comparisons in regulation across the phylogenetic tree, as we have demonstrated with *E. coli*.

## Methods

### RNA extraction and library preparation

The *P. aeruginosa* PAO1 and K2733 (PAO1*ΔmexB*) strains were used in this study. We extracted RNA samples for 25 unique conditions including different media types (M9, CAMHB, LB, RPMI+10%LB), treatment with oxidative stress (Paraquat), iron starvation (DPD), osmotics stress (NaCl), low pH, various carbon sources (succinate, glycerol, pyruvate, fructose, sucrose, N-acetyl glucosamine), micronutrients (copper, iron, zinc, sodium hypochlorite), knockouts, and many more. All conditions were collected in biological duplicates and untreated controls were also collected for each set to rule out the possibility of the batch effect.

In brief, strains were grown overnight at 37°C, with rolling, in appropriate media types for the testing condition of choice. Overnight cultures were then diluted to a starting OD_600_ of ~0.01 and grown at 37°C, with stirring. Once cultures reached the desired OD_600_ of 0.4, 2 mL cultures were immediately added to centrifuge tubes containing 4 mL RNAprotect Bacteria Reagent (Qiagen), vortexed for 5 seconds and incubated at room temperature for 5 min. Samples were then centrifuged for 10 minutes at 5000xg and the supernatant was removed prior to storage at −80°C until further processing. In conditions involving antibiotic treatment, when the bacterial culture had reached an OD_600_ of ~0.2, antibiotics were added at 2X or 5X their MIC in the appropriate media type and allowed to incubate at 37°C, with stirring, for an additional hour prior to sample collection.

Total RNA was isolated and purified using a Zymo Research Quick-RNA Fungal/Bacterial Microprep Kit from frozen cell pellets previously harvested using Qiagen RNAprotect Bacteria Reagent according to the manufacturers’ protocols. Ribosomal RNA was removed from 1 ug Total RNA with the use of a thermostable RNase H (Hybridase) and short DNA oligos complementary to the ribosomal RNA, performed at 65 degrees C to prevent non-specific degradation of mRNA. The resulting rRNA-subtracted RNA was made into libraries with a KAPA RNA HyperPrep kit incorporating short Y-adapters and barcoded PCR primers. The libraries were quantified with a fluorescent assay (dsDNA AccuGreen quantitation kit, Biotium) and checked for proper size distribution and average size with a TapeStation (D1000 Tape, Agilent). Library pools were then assembled and a 1X SPRI bead cleanup performed to remove traces of carryover PCR primers. The final library pool was quantified and run on an Illumina instrument (NextSeq, Novaseq).

### Data acquisition and preprocessing

Apart from the in-house generated data, we also used all the high throughput data available in the literature and obtained from NCBI-SRA. Further, the quality control was done using various filters in multiqc packages like fastqc which includes per_base_sequence_quality, per_tile_sequence_quality, per_base_n_content, and adapter_content ^52^. Further, the aligned reads for the coding sequences (i.e. mRNA) were checked and the too few reads were removed such as to improve the sensitivity. Finally, after the removal of the failed samples, we get 474 samples (97 in-house + 377 literature) for further downstream processing. The raw reads of the RNAseq experiment were mapped on the *P. aeruginosa* reference genome (NC_002516.2) using bowtie2 (*v*2.3.4) ^53^. The ORFs were assigned to the aligned reads through the HTSeq-counts ^54^. Further, the DSeq2 was used to perform the differential expression analysis ^55^, which finally resulted in the transcripts per million (TPM).

### Computing robust independent components

The procedure to calculate the robust independent components was described by McConn *et al* ^7,56^.

In brief, the independent components (ICs) were processed using the decomposition method FastICA available in the Scikit-learn library (*v*0.23.2) ^57^ using the convergence tolerance of 10^−7^, contrast function of log(cosh(x)), 100 iterations, and parallel search. All the components were calculated such as to reconstruct 99% variance during Principal component analysis.

After the 100 iterations of the ICs were clustered using the DBSCAN algorithm with the minimum size of 50 and epsilon 0.1 ^58^. Further, to confirm the reproducibility of the component of each cluster, all the signs of the clusters were inverted in order to get the highest weighted gene with positive sign. The centroids of each cluster were set to the weighting of the ICs.

Further, to confirm the robustness of the components, dimensionality reduction was done. The overall process was repeated multiple times, from dimensions 10 to 420 with the step size of 20. The optimal dimensionality was extracted by comparing the number of ICs with single genes to that of the number of ICs with the largest dimension (PCC 0.70). Finally, the number of dimensions were selected where the number of non-single ICs was equal to the number of final components in that dimension.

### Determination of the gene coefficient threshold

Each gene of 104 components of the M matrix contains a large amount of the values which are near to zeros. Further, the threshold for differentiating the near zero values of the genes to the other contributing values was set in order to identify the set of genes for each iModulon by using the D’Agostino K^2^ test ^59^. The D’Agostino K^2^ statistic test combines the skewness and Kurtosis of the normally distributed components to determine their gaussianity. All genes with the highest absolute weights were removed and the removal process will continue until the K^2^ statistics falls below the threshold. The sensitivity analysis was used to calculate the threshold between the computed iModulons and the regions reported in literature, RegPrecise ^15^, and Pseudomonas genome database ^60^. Further, the threshold of 320 (range) was set to perform the D’Agostino K^2^ test for computing the iModulons through Fisher’s exact test. The precise and recall of all the overlapped iModulons were calculated. Thus, the threshold was chosen which led to the highest harmonic mean (F1) between the recall and precision as described in Sastry *et al* ^4^.

### Regulator enrichment

The transcriptional regulatory network (TRN) was annotated using the information from the RegRrecise^15^, Pseudomonas genome db^60^, and literature. The iModulons with coding RNAs were included in the annotation while the non-coding RNAs were excluded. However, in case of the *P. aeruginosa* very few information of the TRN were available in RegPrecise and Pseudomonas genome db. For the unannotated iModulons we use the Gene Ontology ^61^ and KEGG pathway ^62^ annotations using the precision, recall, *p-value*, and *q-values*. Furthermore, the remaining unannotated iModulons were manually annotated from literature.

### Differential activation analysis

We calculated the difference in the iModulons activities between two or more conditions by fiting the log-normal distribution to the difference. The statistical significance was checked by calculating the difference between the absolute values of the activities among the iModulons, and further confirmed by the *p*-values. However, the *p*-values were adjusted using the Banjamini-Hochberg correction. While calculating the differential activation among the iModulons, the difference with >5 was considered significant.

### Characterizing functionally correlated iModulons

We used the Pearson’s correlation matrix to identify the biologically similar iModulons. Furthermore, Agglomerative (hierarchical) clustering is used under the hood. Thus, a distance threshold for defining “flat” clusters from the hierarchical structure must be determined. By default, this distance threshold is automatically calculated using a sensitivity analysis. Different distance thresholds (this value is between 0 and 1) are tried, and the resulting clustering is assessed using a silhouette score, which is a measure of how separate the clusters are. The distance threshold yielding the maximum silhouette score is automatically chosen.

### Prediction of the biosynthetic gene clusters

We used the antiSMASH algorithm to predict the BGCs in the *P. aeruginosa* ^20^. While using the anti-SMASH software, we used the *P. aeruginosa* (NC_002516.2) with the ‘relaxed’ detection strictness. The antiSMASH algorithm predicts different types of the BGCs like NRPS, PKS, RiPP, redox-cofactors, and many more. Apart from the predicted BGCs, antiSMASH also provides the gene ontology annotations for the BGCs components.

### Generating iModulonDB Dashboards

iModulonDB dashboards were generated using the PyModulon package ^4,12^; the pipeline for doing so can be found at https://pymodulon.readthedocs.io/en/latest/tutorials/creating_an_imodulondb_dashboard.html. Where applicable, we provide links to gene information in Pseudomonas.com ^60^.

### Data availability

All the in-house generated sequences were deposited in the NCBI-Sequence Read Archive database. The accession number of the deposited reads is provided in the **Supplementary Table S7**. While the X, M and A matrices are available in the **Supplementary Table S4, S5, and S6** and GitHub (https://github.com/akanksha-r/modulome_paeru1.0). Each gene and iModulon have interactive, searchable dashboards on iModulonDB.org, and data can also be downloaded from there.

### Code availability

The customized code for the ICA analysis is provided on GitHub (https://github.com/akanksha-r/modulome_paeru1.0). While various files including the X, M, A matrices, TRN regulator file, gene annotated files, gene ontology and kegg pathway annotation files are available on GitHub (https://github.com/akanksha-r/modulome_paeru1.0).

## Acknowledgements

We are grateful to Omkar Satyavan Mohite for the informative discussion on biosynthetic gene cluster analysis. We thank Marc Abrams for reviewing the manuscript and providing constructive suggestions. This work was supported by NIH Grant U01 AI124316 and Novo Nordisk Foundation Grant NNF10CC1016517.

