## Supplementary Figure for "Machine Learning of *Pseudomonas aeruginosa* transcriptomes identifies independently modulated sets of genes associated with known transcriptional regulators"

**Author Affiliations**

**Supplementary Figures**

***Supplementary Figure S1****. Overview of the RNAseq data of Pseudomonas aeruginosa used in the study a) A clustermap shows the global correlations between one sample and all others along with the hierarchical clustering to identify specific clusters. b) Schematic representation of the definitions of regulon recall and iModulon recall. c) Principal component plot shows the diversity of the samples used in the study. The color represents different projects used in the current study*

***Supplementary Figure S2****. Overview of the biosynthetic gene clusters (BGCs) of Pseudomonas aeruginosa predicted from our ICA based analysis. a) Genomic location of the 14 BGCs predicted from the antiSMASH software. b) structure of 11 BGCs defined by our ICA based pipeline*

***Supplementary Figure S3****. Novel RiPP iModulon. a) Scatter plot showing the gene weights of the novel RiPP iModulon, the color depicts the COG categories. b) Activity plot of the conditions expressed in RiPP iModulon in the Paeru_PRECISE*

***Supplementary Figure S4****. Graphical representation of the iModulons predicted and mapped as carbon and amino acid metabolism pathways. The carbon metabolism pathways are ED pathway, glycolysis, and peripheral pathways. While the amino acid metabolism pathways includes the branched chain amino acid (BCAA), aromatic amino acid (AAA), histidine utilization (HUT), arginine decarboxylase (ADC), arginine succinyltransferase (AST), arginine deiminase (ADI), and L-hydroxyproline (LHP) pathways*

***Supplementary Figure S5****. iModulons related to the Carbon metabolism and the Amino Acid/Nucleotide metabolism. a) Activity plot of the conditions expressed in GlpR iModulons in the Paeru_PRECISE. b) Scatter plot showing the correlation between the BCAA pathways iModulons i.e. LiuR and PhhR with the PCC of 0.50.*

***Supplementary Figure S6****. Activity clustering of the iModulons among the P. aeruginosa a) Iron acquisition cluster with iModulons like FpvR, PvdS, Uncharacterized-13, PchR, and FoxR grouped with silhouette score of 0.51. b) The scatter plot showing correlation between the FoxR and PvdS iModulons with PCC of 0.67. Both the iModulons show high activity in the EDTA and planktonic form of growth in P. aeruginosa. c) Scatter plot showing the gene weights between the Uncharacterized-13 and the PvdS iModulons. The red colored genes are common between both iModulons. d) Scatter plot showing correlation between the RpoS-2 iModulon activity and the rpoS gene expression with the Pearson's correlation coefficient of 0.61.*

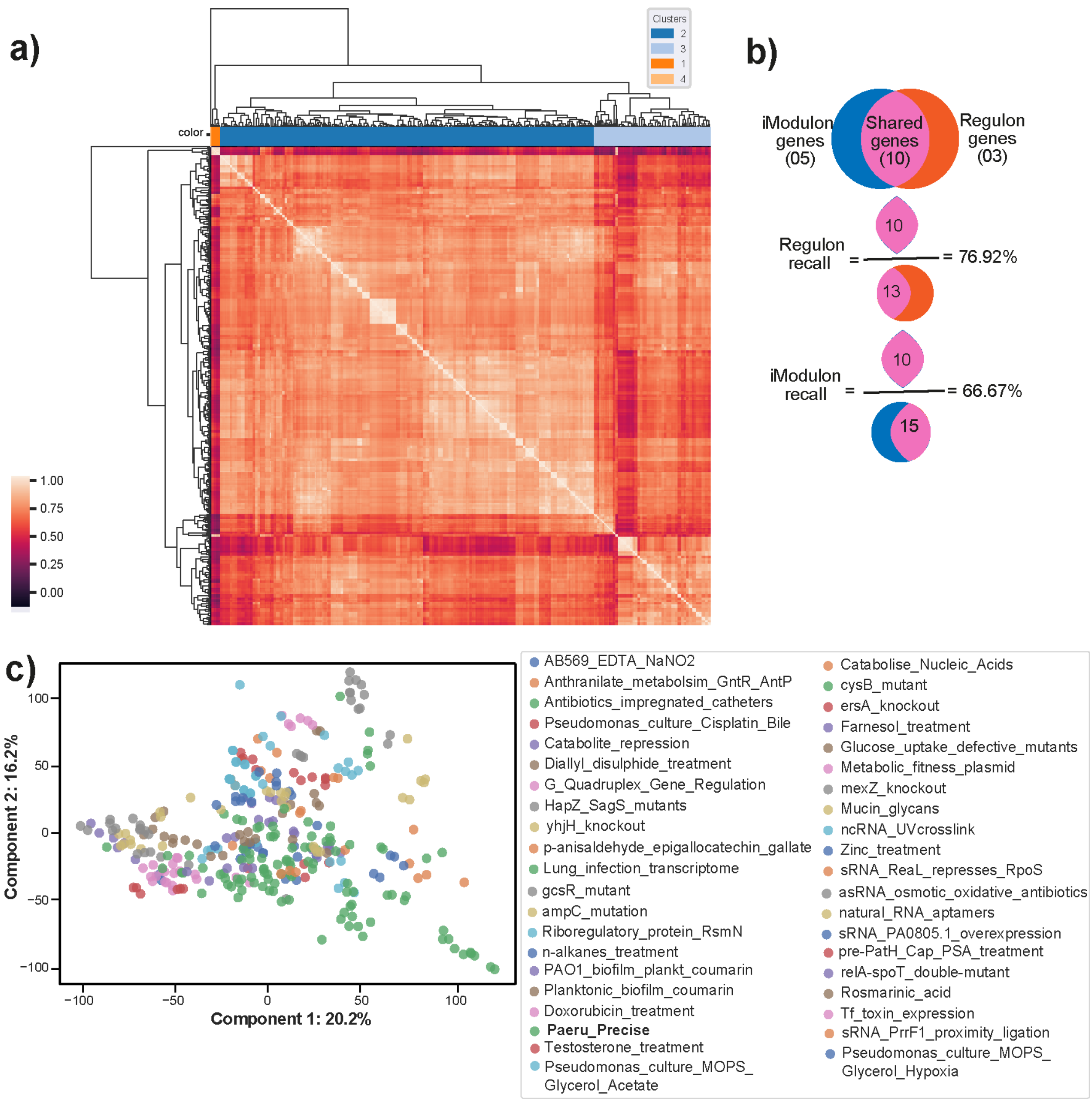


***Supplementary Figure S1**. Overview of the RNAseq data of Pseudomonas aeruginosa used in the study a) A clustermap shows the global correlations between one sample and all others along with the hierarchical clustering to identify specific clusters. b) Schematic representation of the definitions of regulon recall and iModulon recall. c) Principal component plot shows the diversity of the samples used in the study. The color represents different projects used in the current study.*

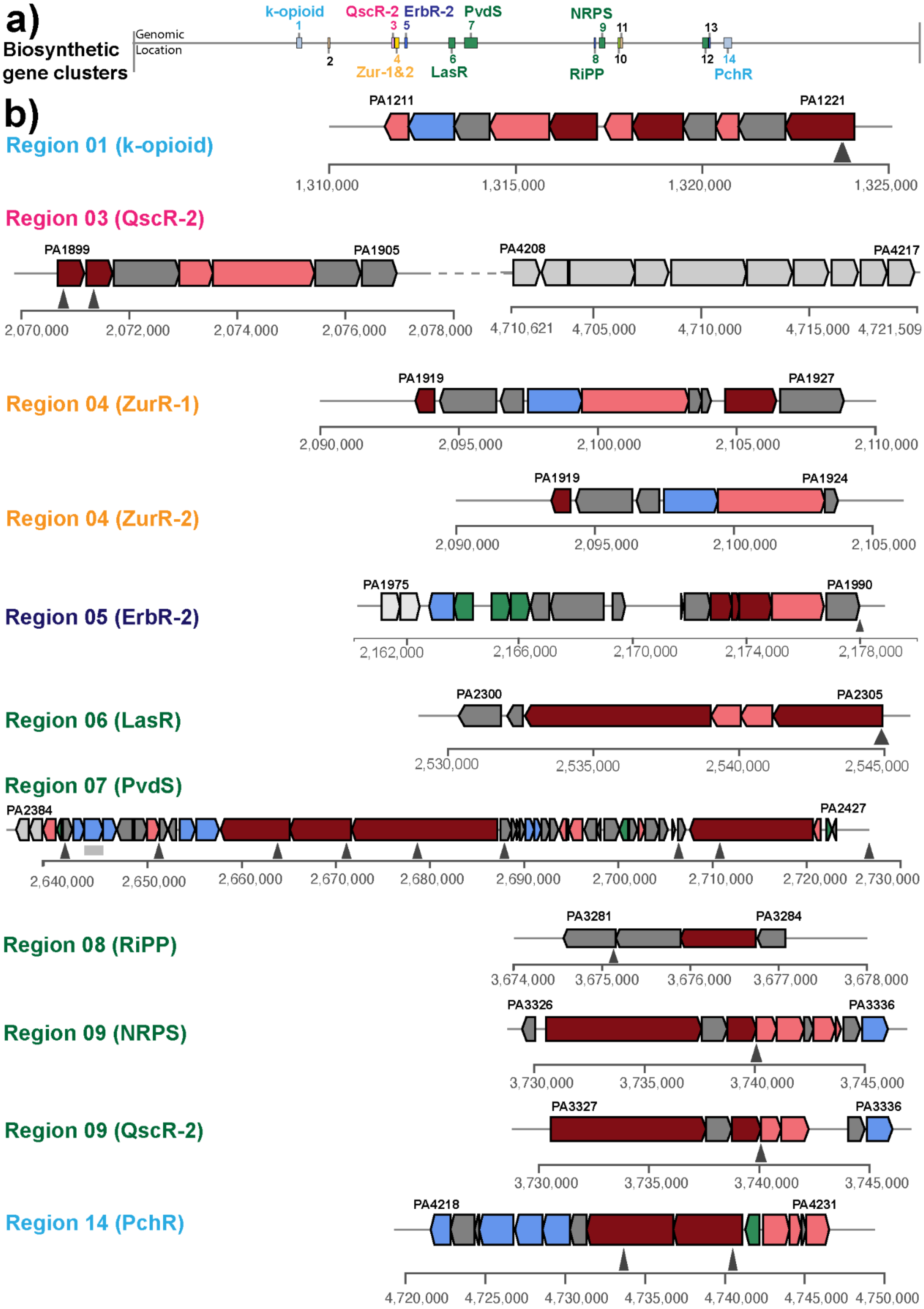
a)
Biosynthetic gene clusters
Genomic Location
k-opioid
QscR-2
ErbR-2
PvdS
NRPS
Zur-1&2
LasR
RiPP
PchR
b)
Region 01 (k-opioid)
PA1211
PA1221
1,310,000
1,315,000
1,320,000
1,325,000
Region 03 (QscR-2)
PA1899
PA1905
PA4208
PA4217
2,070,000
2,072,000
2,074,000
2,076,000
2,078,000
4,710,621
4,705,000
4,710,000
4,715,000
4,721,509
Region 04 (ZurR-1)
PA1919
PA1927
2,090,000
2,095,000
2,100,000
2,105,000
2,110,000
Region 04 (ZurR-2)
PA1919
PA1924
2,090,000
2,095,000
2,100,000
2,105,000
Region 05 (ErbR-2)
PA1975
PA1990
2,162,000
2,166,000
2,170,000
2,174,000
2,178,000
Region 06 (LasR)
PA2300
PA2305
2,530,000
2,535,000
2,540,000
2,545,000
Region 07 (PvdS)
PA2384
PA2427
2,640,000
2,650,000
2,660,000
2,670,000
2,680,000
2,690,000
2,700,000
2,710,000
2,720,000
2,730,000
Region 08 (RiPP)
PA3281
PA3284
3,674,000
3,675,000
3,676,000
3,677,000
3,678,000
Region 09 (NRPS)
PA3326
PA3336
3,730,000
3,735,000
3,740,000
3,745,000
Region 09 (QscR-2)
PA3327
PA3336
3,730,000
3,735,000
3,740,000
3,745,000
Region 14 (PchR)
PA4218
PA4231
4,720,000
4,725,000
4,730,000
4,735,000
4,740,000
4,745,000
4,750,000

***Supplementary Figure S2****. Overview of the biosynthetic gene clusters (BGCs) of Pseudomonas aeruginosa predicted from our ICA based analysis. a) Genomic location of the 14 BGCs predicted from the antiSMASH software. b) structure of 11 BGCs defined by our ICA based pipeline*

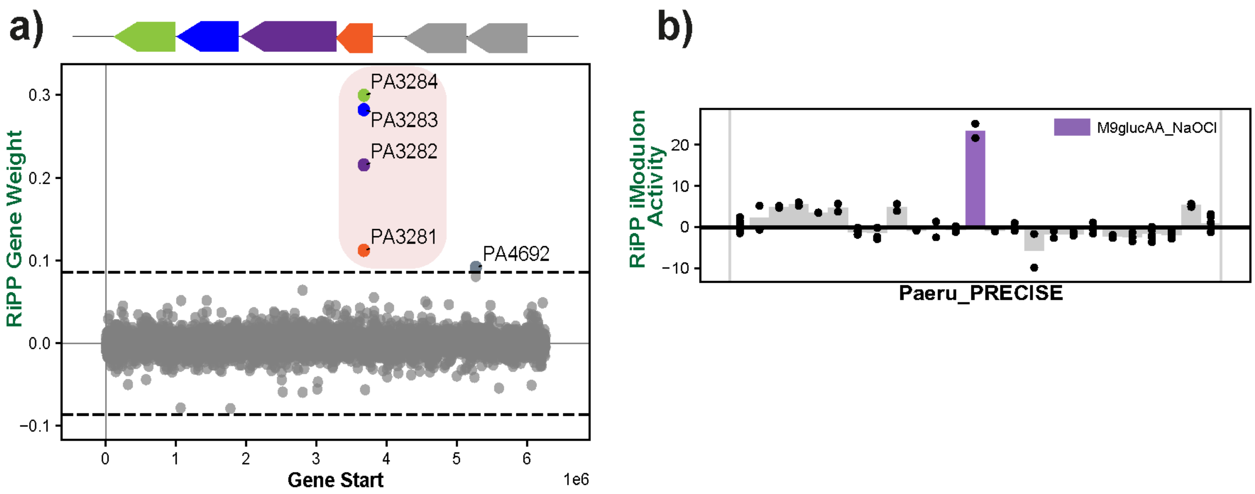


***Supplementary Figure S3****. a) Scatter plot showing the gene weights of the novel RiPP iModulon, the color depicts the COG categories. b) Activity plot of the conditions expressed in RiPP iModulon in the Paeru_PRECISE*

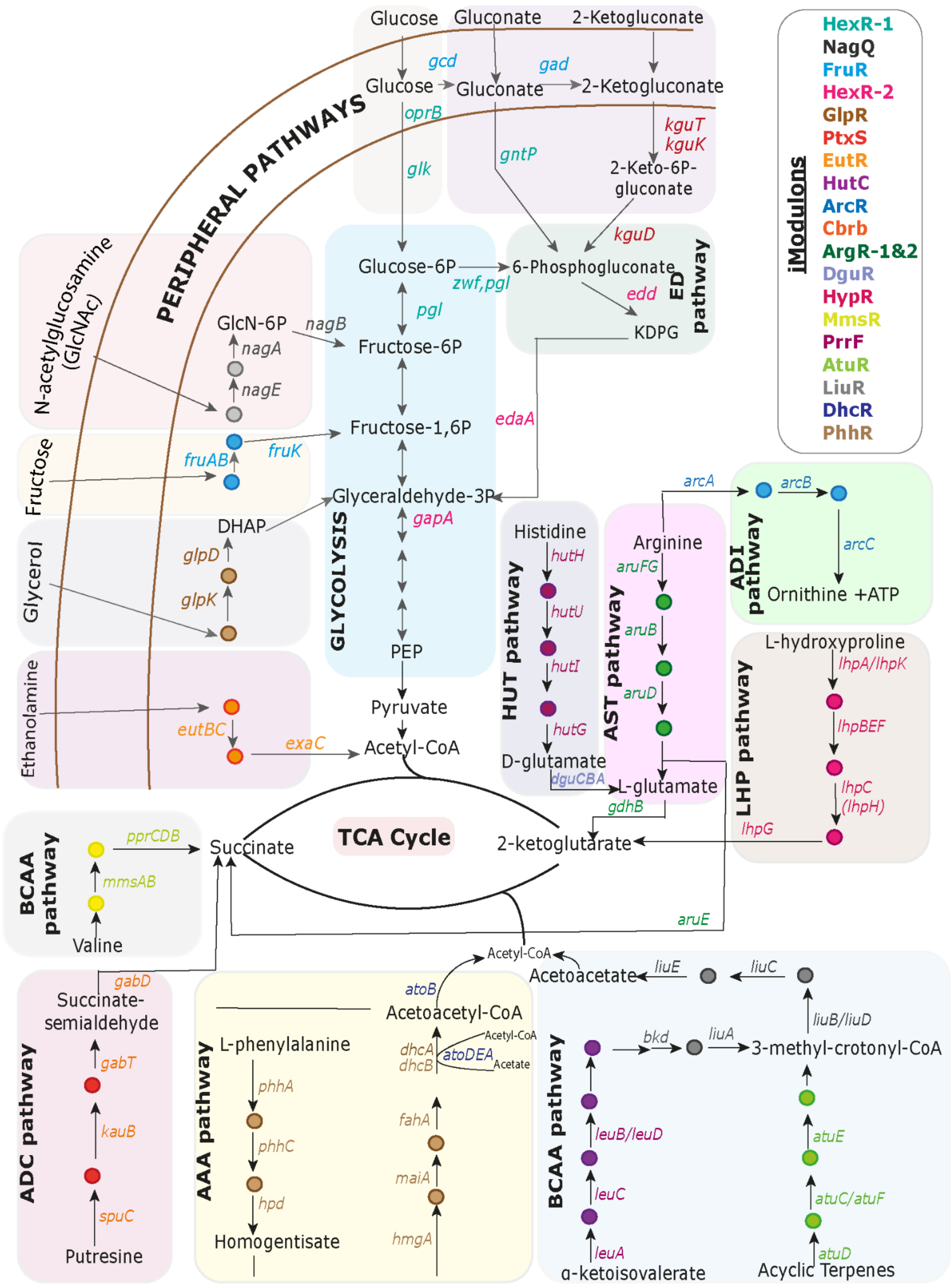

PERIPHERAL PATHWAYS
Glucose
Gluconate
2-Ketogluconate
gcd
gad
oprB
glk
gntP
kguT
kguK
2-Keto-6P-gluconate
kguD
Glucose-6P
6-Phosphogluconate
zwf,pgl
pgl
edd
KDPG
ED pathway
N-acetylglucosamine (GlcNAc)
GlcN-6P
nagB
nagA
nagE
Fructose-6P
Fructose-1,6P
edaA
Fructose
fruAB
fruK
Glyceraldehyde-3P
gapA
DHAP
Glycerol
glpD
glpK
GLYCOLYSIS
PEP
Ethanolamine
eutBC
exaC
Pyruvate
Acetyl-CoA
Histidine
hutH
hutU
hutI
hutG
D-glutamate
dguCBA
HUT pathway
Arginine
aruFG
aruB
aruD
AST pathway
L-glutamate
gdhB
arcA
arcB
arcC
Ornithine +ATP
ADI pathway
L-hydroxyproline
lhpA/lhpK
lhpBEF
lhpC (lhpH)
lhpG
LHP pathway
TCA Cycle
Succinate
2-ketoglutarate
aruE
BCAA pathway
pprCDB
mmsAB
Valine
gabD
Succinate-semialdehyde
gabT
kauB
spuC
Putresine
ADC pathway
Acetyl-CoA
atoB
Acetoacetyl-CoA
Acetoacetate
liuE
liuC
liuB/liuD
dhcA
dhcB
atoDEA
Acetyl-CoA
Acetate
L-phenylalanine
phhA
phhC
hpd
Homogentisate
fahA
maiA
hmgA
AAA pathway
bkd
liuA
3-methyl-crotonyl-CoA
leuB/leuD
leuC
leuA
α-ketoisovalerate
atuE
atuC/atuF
atuD
Acyclic Terpenes
BCAA pathway
iModulons
HexR-1
NagQ
FruR
HexR-2
GlpR
PtxS
EutR
HutC
ArcR
Cbrb
ArgR-1&2
DguR
HypR
MmsR
PrrF
AtuR
LiuR
DhcR
PhhR

***Supplementary Figure S4****. Graphical representation of the iModulons predicted and mapped as carbon and amino acid metabolism pathways. The carbon metabolism pathways are ED pathway, glycolysis, and peripheral pathways. While the amino acid metabolism pathways includes the branched chain amino acid (BCAA), aromatic amino acid (AAA), histidine utilization (HUT), arginine decarboxylase (ADC), arginine succinyltransferase (AST), arginine deiminase (ADI), and L-hydroxyproline (LHP) pathways*

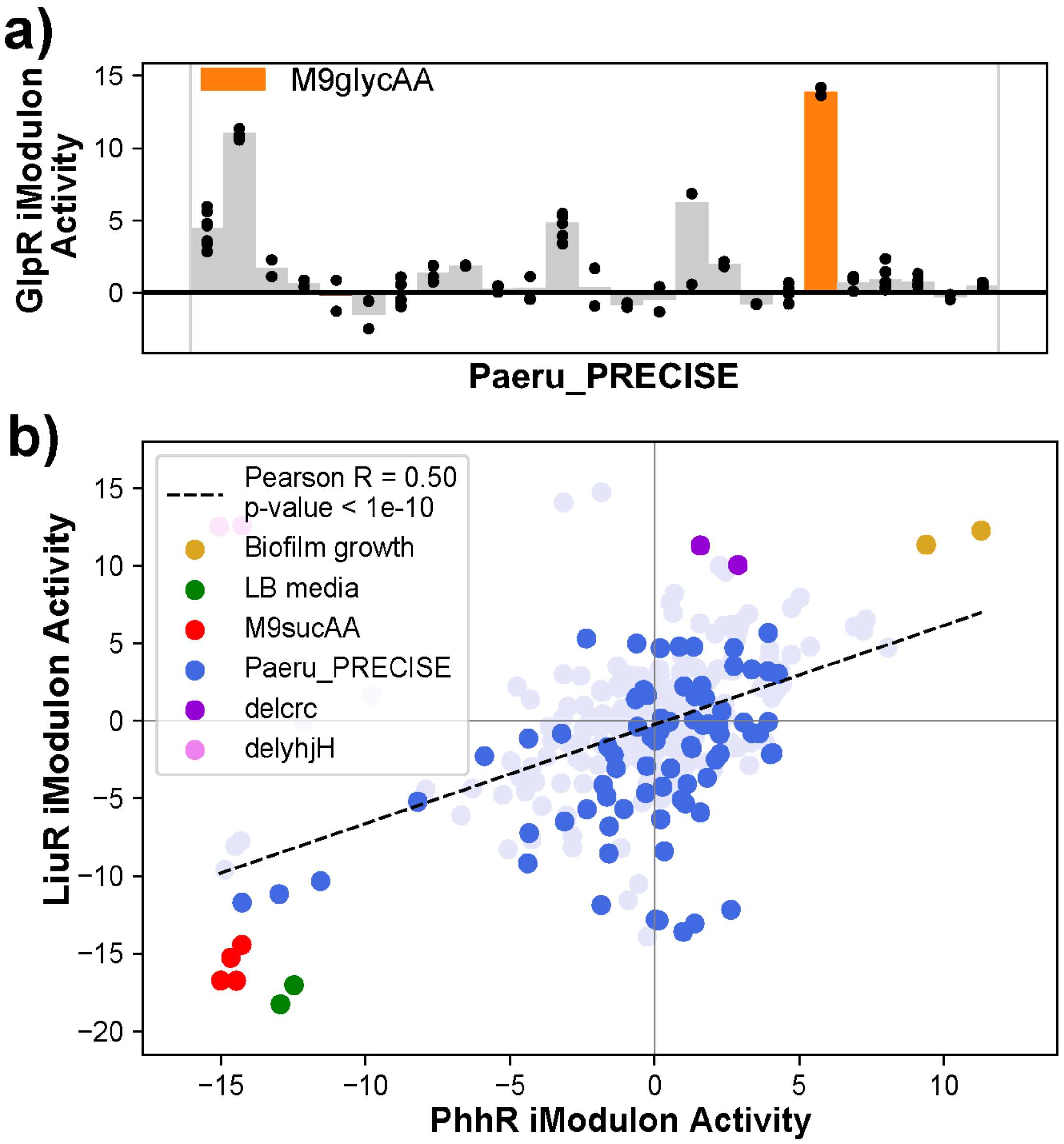


***Supplementary Figure S5****. iModulons related to the Carbon metabolism and the Amino Acid/Nucleotide metabolism. a) Activity plot of the conditions expressed in GlpR iModulons in the Paeru_PRECISE. b) Scatter plot showing the correlation between the BCAA pathways iModulons i.e. LiuR and PhhR with the PCC of 0.50.*

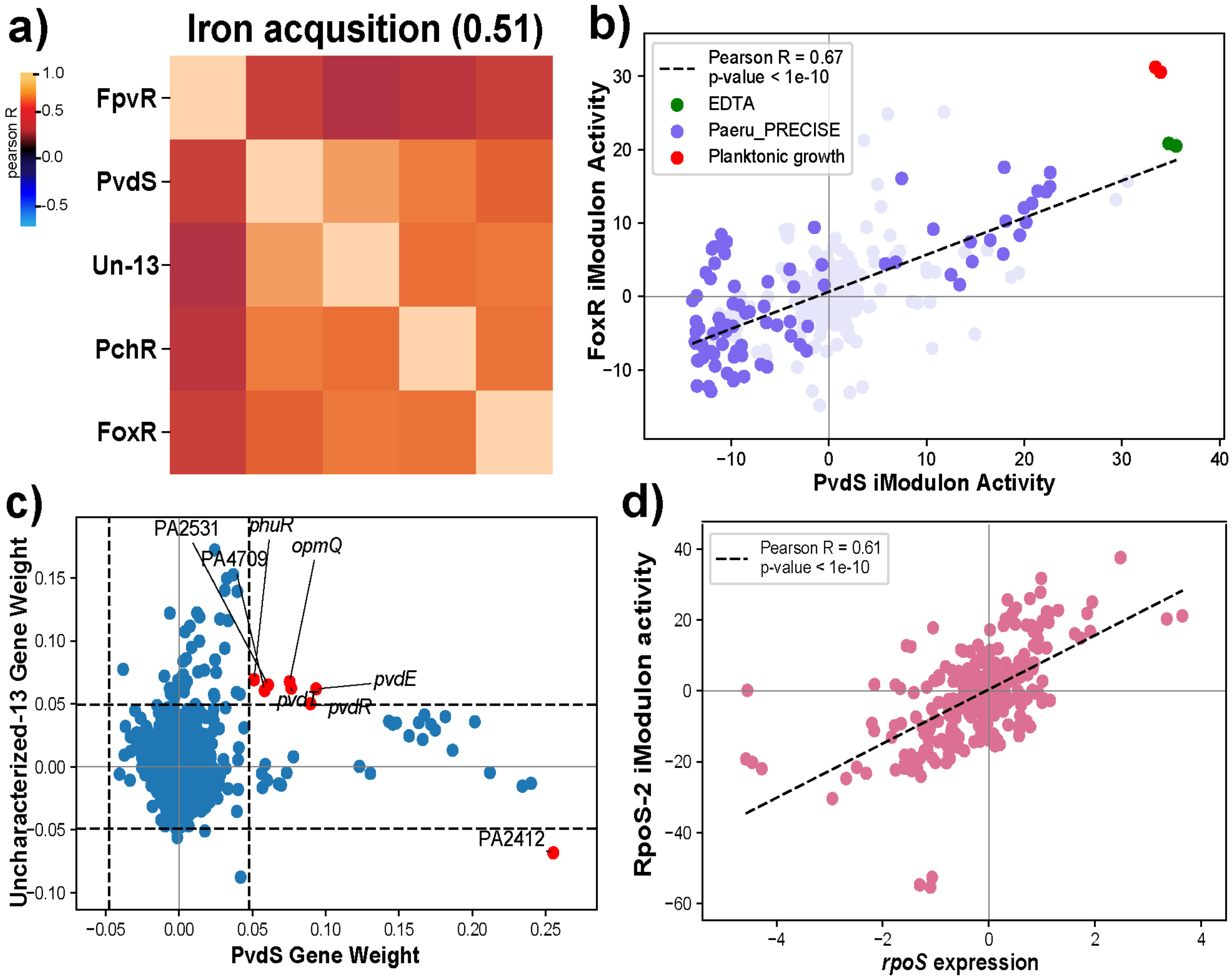


***Supplementary Figure S6****. Activity clustering of the iModulons among the P. aeruginosa a) Iron acquisition cluster with iModulons like FpvR, PvdS, Uncharacterized-13, PchR, and FoxR grouped with silhouette score of 0.51. b) The scatter plot showing correlation between the FoxR and PvdS iModulons with PCC of 0.67. Both the iModulons show high activity in the EDTA and planktonic form of growth in P. aeruginosa. c) Scatter plot showing the gene weights between the Uncharacterized-13 and the PvdS iModulons. The red colored genes are common between both iModulons. d) Scatter plot showing correlation between the RpoS-2 iModulon activity and the rpoS gene expression with the Pearson's correlation coefficient of 0.61.*
